# Excess neonatal testosterone causes male-specific social and fear memory deficits in wild-type mice

**DOI:** 10.1101/2023.10.18.562939

**Authors:** Pravda Quiñones-Labernik, Kelsey L Blocklinger, Matthew R Bruce, Sarah L Ferri

## Abstract

Neurodevelopmental disorders disproportionately affect males compared to females. The biological mechanisms of this male susceptibility or female protection have not been identified. There is evidence that fetal/neonatal gonadal hormones, which play a pivotal role in many aspects of development, may contribute. Here, we investigate the effects of excess testosterone during a critical period of sex-specific brain organization on social approach and fear learning behaviors in C57BL/6J wild-type mice. Male, but not female, mice treated with testosterone on the day of birth (PN0) exhibited decreased social approach as juveniles and decreased contextual fear memory as adults, compared to vehicle-treated controls. These deficits were not driven by anxiety-like behavior or changes in locomotion or body weight. Mice treated with the same dose of testosterone on postnatal day 18 (PN18), which is outside of the critical period of brain masculinization, did not demonstrate impairments compared to the vehicle group. These findings indicate that excess testosterone during a critical period of early development, but not shortly after, induces long-term deficits relevant to the male sex bias in neurodevelopmental disorders.

**Significance statement:** Excess testosterone during a critical period of sex-specific brain organization, results in male-specific social and cognitive deficits in mice while testosterone treatment outside of this developmental window did not alter behavior. This time-sensitive, brief hormonal dysregulation induces long-term changes, and may be involved in the male sex bias in neurodevelopmental disorders.

## Introduction

Neurodevelopmental and neuropsychiatric conditions are prevalent and can be difficult to treat, emotionally, physically, and financially taxing to affected individuals, their families, and communities (Zablotsky et al., 2019). Therefore, it is crucial to determine developmental processes impacting vulnerability to these disorders, a number of which affect males and females divergently. For example, Autism Spectrum Disorder (ASD) and Attention Deficit Hyperactivity Disorder (ADHD) affect males at a ∼4 and ∼2 to 1 ratio, respectively (Maenner et al., 2023; Yang et al., 2022). Other factors such as age of onset, severity, presentation of symptoms, and response to treatment can also affect males and females differently. The mechanisms underlying these sex differences are not fully understood, but elucidating them would significantly improve risk assessment, early intervention, and targeted treatments.

One of the earliest developmental processes that may be involved in sex-sensitive susceptibility is gonadal hormone exposure during the prenatal and neonatal periods. Human male testosterone production surges mid-gestation and shortly after birth, while male rodents have highest levels several days before and after birth. Females of those species are exposed to significantly lower levels of gonadal hormones during early development (Gillies and McArthur, 2010a; McCarthy, 2011). In rodents, many important and irreversible sex hormone-mediated effects on brain development are complete by postnatal day 10 (Davis et al., 1996; McCarthy et al., 2017). After, sex hormones have more transient effects. It is during the critical period that hormonal dysregulation may cause sex-specific vulnerabilities to neurodevelopmental deficits. There is evidence that males exposed to increased levels of steroidogenic factors in amniotic fluid, likely indicating fetal origin, are more likely to be diagnosed with ASD (Baron-Cohen et al., 2015). Amniotic fluid is usually collected mid-gestation, near the peak production of testosterone in males (Van De Beek et al., 2004). Amniotic testosterone level is correlated with ASD-related traits in children regardless of diagnosis (Auyeung et al., 2012, 2010; Chapman et al., 2006; Knickmeyer and Baron-Cohen, 2006; Lutchmaya et al., 2002a, 2002b). Maternal testosterone in the second trimester is associated with increased autism-relevant behaviors in male children (Firestein et al., 2022). There are a number of conditions in which a fetus may be exposed to excess testosterone derived from the mother or placenta (Baron-Cohen et al., 2015). Congenital adrenal hyperplasia (CAH), Polycystic Ovarian Syndrome (PCOS), pre-eclampsia, maternal diabetes, and maternal stress are conditions with increased androgen levels and increased offspring risk for neurodevelopmental disorders (Codner et al., 2012; Gumusoglu et al., 2020; Kumar et al., 2018; Lai et al., 2011; Li et al., 2023; Meng et al., 2023; Rowland and Wilson, 2021; Xiang et al., 2018). Some studies using digit ratio (2D:4D) or facial landmark masculinity as indirect measures of fetal testosterone exposure have found evidence of higher levels in children later diagnosed with ASD compared to those of typically developing children (De Bruin et al., 2006; Manning et al., 2001; McKenna et al., 2021; Milne et al., 2006).

Many neuropsychiatric conditions present with changes in emotional response and impairments in approach and avoidance behaviors, including sociability and fear. Social behavior is disrupted in ASD, ADHD, schizophrenia, PTSD, psychopathy, and others (Frye, 2018; Homberg et al., 2016; Kennedy and Adolphs, 2012; Nijmeijer et al., 2008; Porcelli et al., 2019; Scoglio et al., 2022). Many social behaviors exhibit sex differences and are sensitive to gonadal hormones and endocrine disruptors (Bell, 2018). Acquisition of conditioned fear and fear extinction is impaired in psychopathy, PTSD, anxiety, and other, disorders (VanElzakker et al., 2014; Veit et al., 2013). The ability to interpret and transmit both social and fear cues are critically important for physical and psychological well-being across many species. In the present study we sought to determine the effects of brief steroid hormone dysregulation during a critical developmental timepoint on social approach and fear memory behaviors. Rodents are a valuable experimental model in which manipulation is well-controlled in timing, consistent, and causal. We administered a single dose of testosterone that has been shown to masculinize a female rodent brain to male and female C57BL/6J pups on the day of birth, during the critical period of sex-specific brain organization (Davis et al., 1996; Ghahramani et al., 2014; Goel and Bale, 2013; Hisasue et al., 2010; McCarthy et al., 2017; Seney et al., 2012). We tested the mice in the three-chamber social approach assay as juveniles and in contextual fear conditioning as adults (**Fig. 1**). We found that a single administration of testosterone during a critical period of brain development (PN0), but not after (PN18), resulted in male-specific deficits in social approach and contextual fear conditioning that were not driven by changes in body weight or anxiety-like behavior.

**Figure 1.**
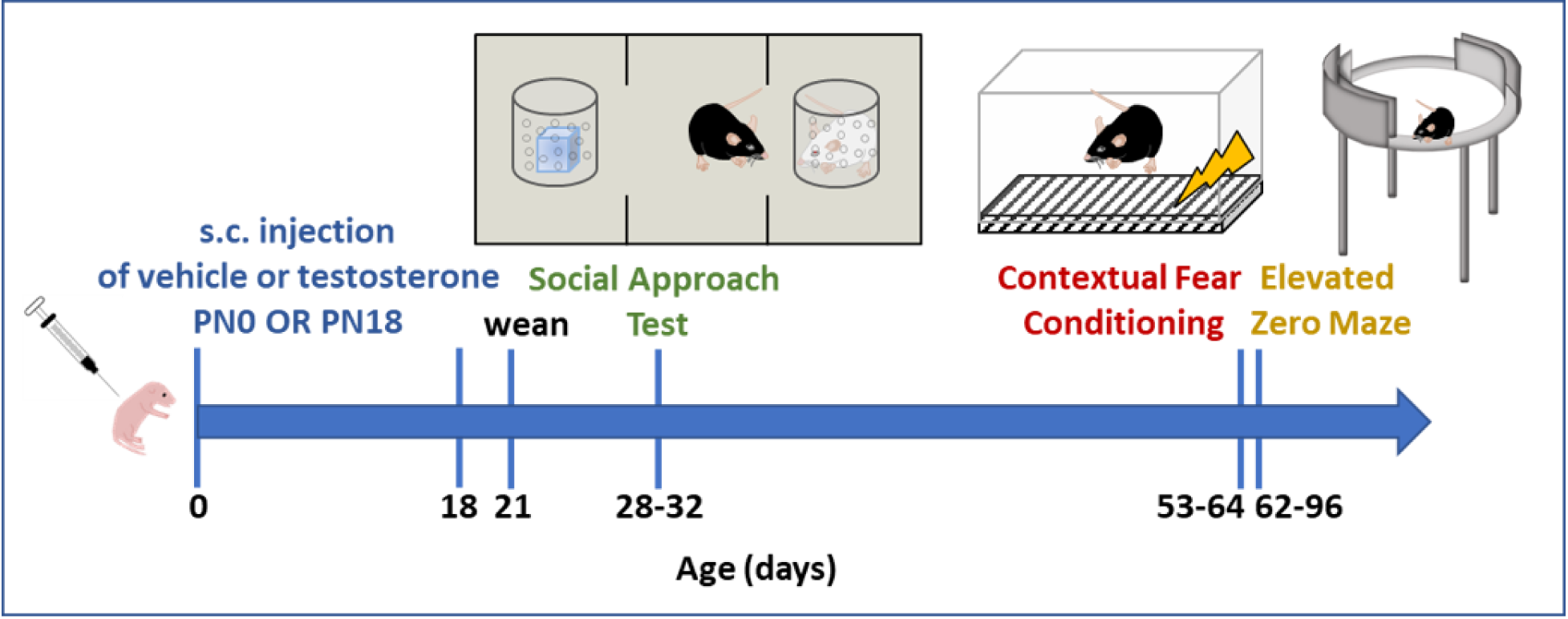
Experimental overview. Pups were injected subcutaneously on the day of birth (postnatal day 0) with testosterone or oil vehicle then remained with the dam and sire until weaning at PN21, after which they were subjected to a weight study (not shown here, PN2-60), social approach test at PN28-32 and contextual fear conditioning at PN53-64, or elevated zero maze at PN62-96. Another group of mice was administered the same dose of testosterone or vehicle on PN18, weaned at PN21, and underwent social approach test at PN28-32 and contextual fear conditioning at PN53-64.

## Materials and Methods

### Animals

Experiments were conducted in accordance with the [Author University] Institutional Animal Care and Use Committee (IACUC) policies and approved protocols, and according to ARRIVE guidelines. Mice were maintained consistent with the Guide for the Care and Use of Laboratory Animals. Mice were housed in a temperature- and humidity-controlled environment (22 °C and 55 ± 5%, respectively). All animals were housed on a 12-hour light/dark cycle (lights-on at 9:00 am). Food and water were available *ad libitum*. C57BL/6J (B6) mice were obtained from The Jackson Laboratories (#000664) to establish breeding cages, which contained one dam and one sire, which remained with the pups until they were weaned. Litters were randomly divided into two experimental conditions: treatment with testosterone (T) or vehicle (veh; see below), receiving treatment on the day of birth (PN0) or on postnatal day 18 (PN18). On postnatal day 21 mice were weaned and housed in groups of 2-5 same-sex littermates per cage, unless otherwise specified. Gonadectomized A/J mice (#000646) were obtained from The Jackson Laboratories and were used as a social stimulus in the social behavioral assay. A total of 205 mice were used for behavioral testing. Of the 205 mice, 21 were excluded due to inactivity during the testing period, excessive climbing on cylinders, or statistical outliers (>2SD from the mean). Of mice treated on PN0, one cohort was used in weight studies, another to test anxiety-like behavior, and another in the social approach test, with a subset undergoing contextual fear conditioning, further outlined below. Mice treated on PN18 completed social approach and fear memory tests at PN28-32 and 53-64, respectively, except for 7/51 mice underwent only fear conditioning as adults.

### Testosterone Treatment

Testosterone propionate (Sigma; 100µg in 20 µl sesame oil, a dose previously shown to induce brain masculinization in a female rodent) (Ghahramani et al., 2014; Goel and Bale, 2013; Hisasue et al., 2010; Nakachi et al., 2015; Seney et al., 2012) or vehicle (20 µl sesame oil) was administered subcutaneously at one of two time points: to pups on the day of birth (PN0) or on postnatal day 18 (PN18). Each individual mouse within a litter received the same treatment so experimental cages of littermates were all of the same treatment group. Breeding cages were checked 2-4 times per day for litters, and pups were injected usually within 4, but no more than 12, hours of birth. Neonates were separated from the dam for less than two minutes and gently scruffed for the subcutaneous injection. Following treatment, pups were returned to their homecage with the dam and sire until weaning.

### Weight Measurement

In an independent cohort of animals, to avoid repeated handling prior to behavioral assays, weights were measured every two days, beginning on day 2 after birth (PN2), until day 12. Then, weights were recorded on day 21, 30, and 60 to monitor treatment effects.

### Social Approach Test

The Social Approach Test was conducted in PN28-32 mice, in a dimly lit room (<5 lux), using a black Plexiglass arena (10 x 20.5 x 9 in) that had three chambers devoid of top and bottom, which was placed on a clear Plexiglass table over a clean absorbent pad. Identical bottomless and topless clear cylinders were placed at the center of both outer arena chambers. Each clear cylinder featured one end with small breathing holes which facilitated air circulation and enabled visual and olfactory exploration. Flat lids were placed on top of the cylinders and secured with small paperweights. The arena was illuminated from below using infrared light. Testing sessions were recorded from an overheard-positioned camera (Basler Ace GIGE). The behavioral assay consisted of two ten-minute phases: a “Habituation” phase followed by a “Choice” phase. During the habituation phase, the test mouse could freely explore the chamber and empty cylinders.

Following the completion of the habituation phase, a novel-object (Duplo block) was introduced into one cylinder, while a novel social stimulus, a same-sex gonadectomized A/J mouse, was placed in the opposite cylinder. Again, the mouse was able to freely explore for 10 minutes during the choice phase. Distance traveled and duration of sniffing of each cylinder were quantified using Noldus EthoVision XT video tracking software. A preference index (PI) was calculated for each phase: (time spent sniffing social cylinder (empty in habituation or containing novel mouse during choice phase) – time spent sniffing nonsocial cylinder (empty or novel object)) / (total sniffing time). A preference index of 0 indicates no preference for either cylinder (equal sniffing of each) and PI of 1 indicates 100% sniffing of social cylinder in the choice phase (or empty cylinder in the habituation phase).

### Contextual Fear Conditioning

After initial data collection in the social approach test, we decided to test contextual fear conditioning in subsequent mice. Of the 69 animals treated at PN0 used in the social approach test at PN28-32, 44 underwent contextual fear conditioning at PN53-64 to avoid testing during the pubertal period. Of the mice treated at PN18, 44 were tested at PN28-32 for social approach and all were subsequently tested at PN53-64 for fear conditioning plus an additional 8 that did not undergo social testing due to a scheduling issue. Mice were singly housed 4-7 days prior to conditioning and handled 2-3 min each for 3 consecutive days prior to the assay. In our hands, this brief single-housing and handling protocol facilitates learning and avoids cagemate fighting, which we often observe in group-housed mice post-shock. On the day of training, each mouse was placed inside a chamber with electrified metal grid flooring (CleverSys) inside a sound-attenuating box (Med Associates) for a duration of 3 min. During the initial 2min and 28s, the mice were allowed to freely explore the chamber, which served as a “baseline” period. After this time, a single 1.5mA footshock was delivered to the mice for 2s. The mice were removed 30s following the shock. The test session was conducted 24 hr later, during which the mice were placed in the same chamber for a period of 5 min. The Cleversys Freezescan software was utilized to record the freezing behavior of the mice.

### Elevated Zero Maze

A separate cohort of naïve adult male and female mice (PN62-96) were utilized to assess anxiety-like behavior in PN0 testosterone- and vehicle-treated mice using an elevated zero maze (EZM). The elevated zero maze is an elevated ring-shaped runway with two open arms and two opposing closed arms. The open arms are devoid of walls resulting in an exposed environment, while the closed arms are enclosed with walls. The elevated zero maze was positioned beneath a camera (Basler Ace GIGE) in 250 lux lighting conditions. MediaRecorder software was used to record the trials. Mice were placed on a boundary between an open and closed area facing the closed area. Each mouse was given a 5-min trial, during which they were allowed to freely roam the maze. The experimenter positioned themself behind a white curtain throughout the trial. The maze was cleaned with paper towels and 70% ethanol between each trial. Noldus Ethovision XT video tracking software was used to analyze time spent in the open versus closed areas and total distance traveled within the arena.

### Statistical Analysis

Statistical analysis was performed in GraphPad Prism 9. Weight data were analyzed used a repeated measures three-way ANOVA to determine main effects of time, sex, treatment, and interactions. For all other data, a two-way ANOVA was performed with sex and treatment as main effects, and a sex x treatment interaction was also tested. A Tukey post hoc test was used when appropriate. Spearman’s test was used to determine heteroscedasticity and Shapiro-Wilk for test of normality. Log transformation was used in case of violations, however, raw data is shown in the graph for clarity. Significance was set to p<0.05. Bar graphs and error bars represent mean ± SEM and individual data points are shown. Eta squared values were used for effect size estimations.

## Results

### A single testosterone treatment on the day of birth results in male-specific social approach deficits in juveniles

We first aimed to determine whether excess neonatal testosterone would affect social approach behavior. For the preference index (PI) in the habituation phase, a two-way ANOVA revealed no main effect of sex (F_(1,65)_=0.200, p=0.656, ƞ^2^=0.003), treatment (F_(1,65)_=8.17, p=0.370, ƞ^2^=0.012), or sex x treatment interaction (F_(1,65)_=0.034, p=0.855, ƞ^2^=0.0005; **Fig. 2A**). A two-way ANOVA of the PI in the choice phase uncovered no main effect of sex (F_(1,65)_=2.188, p=0.144, ƞ^2^=0.028), a main effect of treatment (F_(1,65)_=5.743, p=0.019, ƞ^2^=0.073), and a statistical trend of a sex x treatment interaction (F_(1,65)_=3.827, p=0.055, ƞ^2^=0.049). A Tukey post hoc test indicated that males treated on PN0 with testosterone (Males + T) had significantly lower PI than Males + Veh, Females + Veh, and Females + T (p=0.013, 0.039, and 0.038, respectively; **Fig. 2B**).

**Figure 2.**
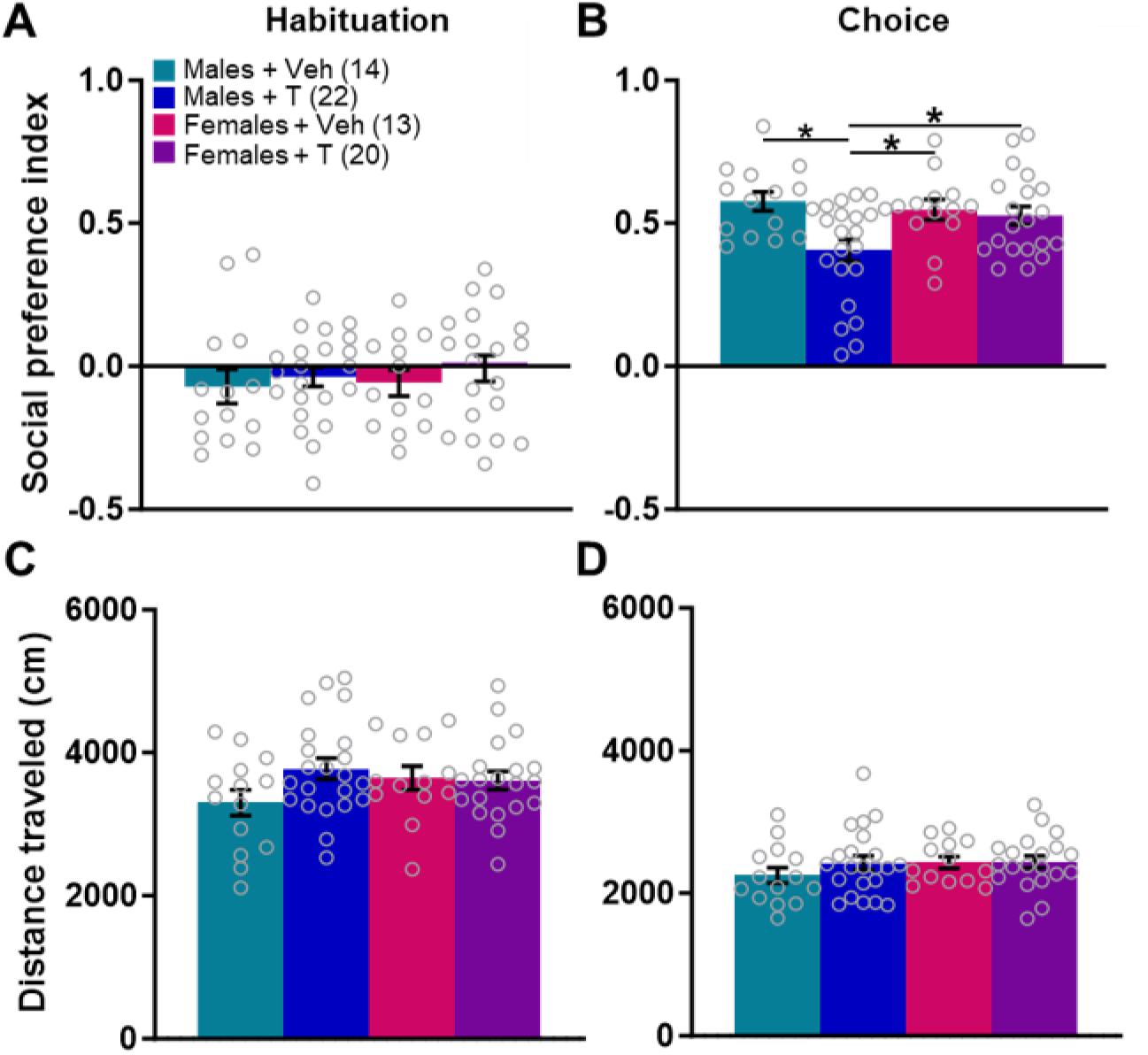
Testosterone administration on the day of birth induces social approach deficits in adolescent males. (A) During a 10 min habituation period of the social approach test, all experimental groups had a similar preference index (PI). (B) Males treated neonatally with testosterone had a significantly lower social preference index than males or females treated with vehicle or females treated with testosterone during the choice phase of the social approach test. There were no differences across groups in distance traveled during the social approach test in either the habituation (C) or choice phase (D).*p<0.05. Bars indicate mean ± SEM.

Neonatal testosterone treatment had no effects on distance traveled during the social approach test. In juvenile male mice (PN28-32) treated with testosterone at PN0, a two-way ANOVA revealed no main effect of sex (F_(1,65)_=0.355, p=0.553, ƞ^2^=0.005) or treatment (F_(1,65)_=2.008, p=0.161, ƞ^2^=0.029), and no significant interaction during the habituation phase (F_(1,65)_=2.655, p=0.108, ƞ^2^=0.038; **Fig. 2C**). Similarly, during the choice phase, there was no main effect of sex (F_(1,65)_=1.029, p=0.314, ƞ^2^=0.015) or treatment (F_(1,65)_=0.823, p=0.368, ƞ^2^=0.012), and no significant sex x treatment interaction (F_(1,65)_=0.697, p=0.407, ƞ^2^=0.010; two-way ANOVA; **Fig. 2D**). Therefore, the decrease in social approach in juvenile males treated neonatally with testosterone was not an effect of altered locomotor behavior.

### A single testosterone treatment on the day of birth results in male-specific contextual fear conditioning deficits in adults

After initial data collection revealed a male-specific social deficit, we used subsequent animals for contextual fear conditioning following social testing, but we wanted to avoid testing them during puberty, which involves many dynamic changes, so we waited until young adulthood.

Therefore, a subset of animals treated with veh or T on PN0 were tested for social approach as juveniles (PN28-32) and then additionally underwent 24-hr fear memory testing as adults (PN53-64). Percent time spent freezing during the pre-shock baseline period was similar across groups; a two-way ANOVA showed no main effect of sex (F_(1,40)_=0.199, p=0.658, ƞ^2^=0.005), treatment (F_(1,40)_=2.067, p=0.158, ƞ^2^=0.047), or sex x treatment interaction (F_(1,40)_=1.372, p=0.248, ƞ^2^=0.031; **Fig. 3A**). A two-way ANOVA of freezing during the 24 h memory test uncovered a significant main effect of sex (F_(1,40)_=4.514, p=0.040, ƞ^2^=0.087), treatment (F_(1,40)_=4.310, p=0.044, ƞ^2^=0.084), and interaction (F_(1,40)_=4.269, p=0.045, ƞ^2^=0.082). A Tukey post hoc test indicated that males treated on PN0 with testosterone (Males + T) had significantly lower freezing than Males + Veh, Females + Veh, and Females + T (p=0.029, 0.034, and 0.035, respectively; **Fig. 3B**).

**Figure 3.**
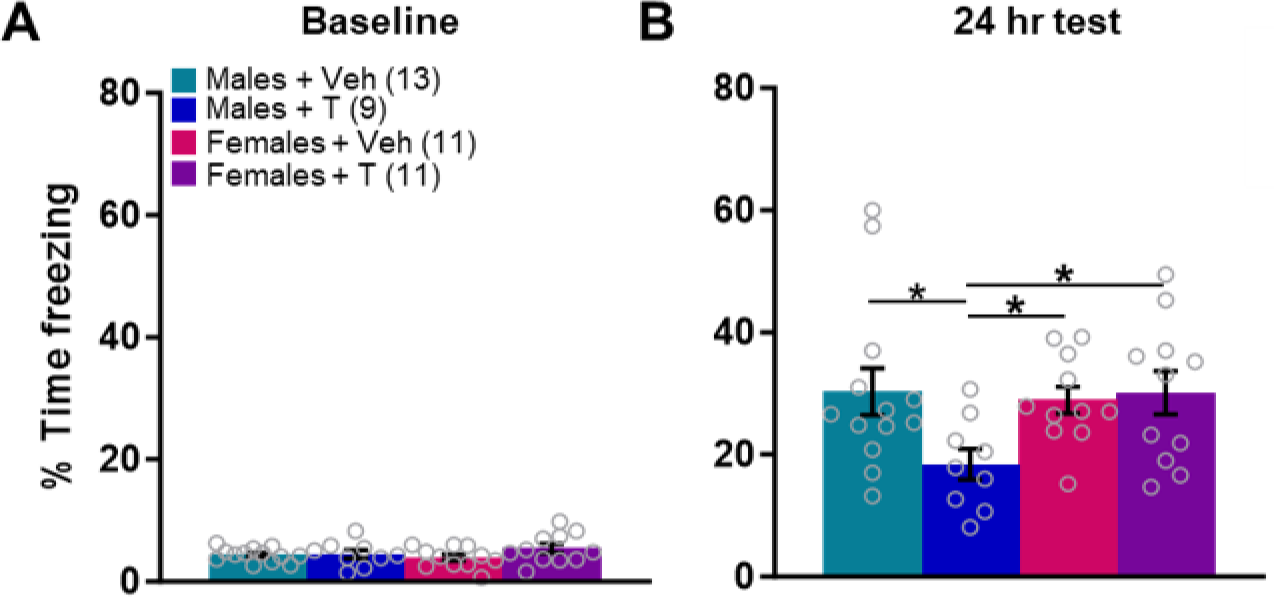
Testosterone administration on the day of birth induces fear memory deficits in adult males. (A) Baseline freezing prior to contextual fear conditioning was similar regardless of sex or treatment. (B) Adult males treated on the day of birth exhibited significantly less freezing during the 24 hr memory test than those treated with veh or females treated with either veh or T. *p<0.05. Bars indicate mean ± SEM.

### Social and fear memory deficits are not due to differences in body weight or anxiety-like behavior

We then used separate groups of animals to determine whether changes in body weight or anxiety may contribute, indicating a general disruption in growth or development that may affect behavior or movement, or decreased overall exploration or increased avoidance. Mice that underwent treatment with neonatal testosterone had similar body weight to those treated with veh on the day of birth (**Fig. 4A**). A RM three-way ANOVA revealed main effects of age (F_(2,60)_=2525, p<0.0001, ƞ^2^=0.886), sex (F_(1,29)_=12.69, p=0.001, ƞ^2^=0.004), and treatment (veh vs T; F_(1,29)_=7.345, p=0.011, ƞ^2^=0.002), and the following significant interactions: age x sex (F_(8,232)_=41.04, p<0.0001, ƞ^2^=0.014), age x treatment (F_(8,232)_=4.967, p<0.0001, ƞ^2^=0.002), and age x sex x treatment (F_(8,232)_=2.556, p<0.011, ƞ^2^=0.0008), but no sex x treatment interaction (F_(1,29)_=0.027, p=0.871, ƞ^2^=7.9e-6). A Tukey post hoc test indicated that at age PN30, males treated with vehicle weighed significantly more than females treated with testosterone (p=0.021), and at PN60, Males + Veh weighed significantly more than both Females + Veh and Females + T (p<0.0001 for both). In summary, most significant differences in weight were due to sex as expected, but there were no significant differences driven by testosterone within sexes at any age.

**Figure 4.**
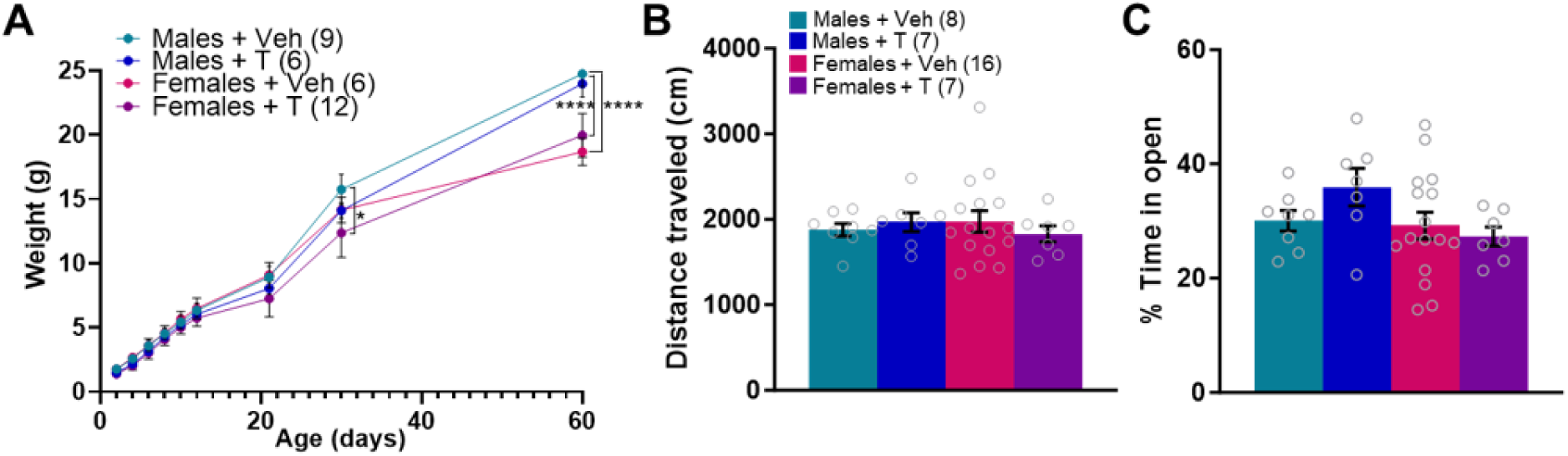
Excess neonatal testosterone-induced effects are not due to changes in weight or anxiety-like behavior. (A) Testosterone administration on the day of birth does not affect body weight. Mice treated at PN0 with testosterone or veh were weighed on PN2, 4, 6, 8, 10, 12, 21, 30, and 60. Most differences in weight were driven by sex. Specifically, at PN30, males treated with vehicle weighed significantly more than females treated with testosterone and at PN60, males treated neonatally with vehicle weighed significantly more than females treated either with vehicle or testosterone on the day of birth. (B) Naïve adult males and females treated neonatally with testosterone traveled similar distances in the elevated zero maze as those treated with veh. (C) Testosterone treatment on the day of birth had no effect on percent time spent in the open arms of the EZM. *p<0.05, ****p<0.0001.

Naïve adult mice treated on the day of birth with testosterone did not show any differences in distance traveled or percent time spent in open arms in the elevated zero maze compared to those treated with vehicle on the day of birth (**Fig 4B** and **4C**). A two-way ANOVA of total distance traveled over the 5-min test revealed no main effect of sex (F_(1,34)_=0.102, p=0.751, ƞ^2^=0.003) or treatment (F_(1,34)_=0.012, p=0.913, ƞ^2^=0.0003) and no interaction between the two (F_(1,34)_=0.623, p=0.435, ƞ^2^=0.018). A two-way ANOVA of percent time spent in the open arms also revealed no differences, with no main effect of sex (F_(1,34)_=3.117, p=0.087, ƞ^2^=0.08) or treatment (F_(1,34)_=0.551, p=0.463, ƞ^2^=0.014) and no interaction between the two (F_(1,34)_=2.090, p=0.158, ƞ^2^=0.054).

### A single testosterone treatment on PN18 does not induce social approach deficits in juveniles

Next, we wanted to determine if the same dose of testosterone treatment given later, outside the reported critical period of brain masculinization (Davis et al., 1996; McCarthy et al., 2017), would cause similar social impairments. For PI during the habituation phase of juvenile mice treated with veh or testosterone on PN18, a two-way ANOVA showed no main effects or interaction (F_(1,39)_=0.010, p=0.922, ƞ^2^=0.0002 for sex; F_(1,39)_=1.280, p=0.265, ƞ^2^=0.031 for treatment; F_(1,39)_=0.586, p=0.449, ƞ^2^=0.014 for interaction; **Fig. 5A**). During the choice phase of the social approach test, all groups had a similar PI; a two-way ANOVA produced no main effects of sex (F_(1,39)_=0.084, p=0.773, ƞ^2^=0.022) or treatment (F_(1,39)_=0.001, p=0.971, ƞ^2^=0.00003), and no interaction of the two (F_(1,39)_=0.120, p=0.731, ƞ^2^=0.003; **Fig. 5B**).

**Figure 5.**
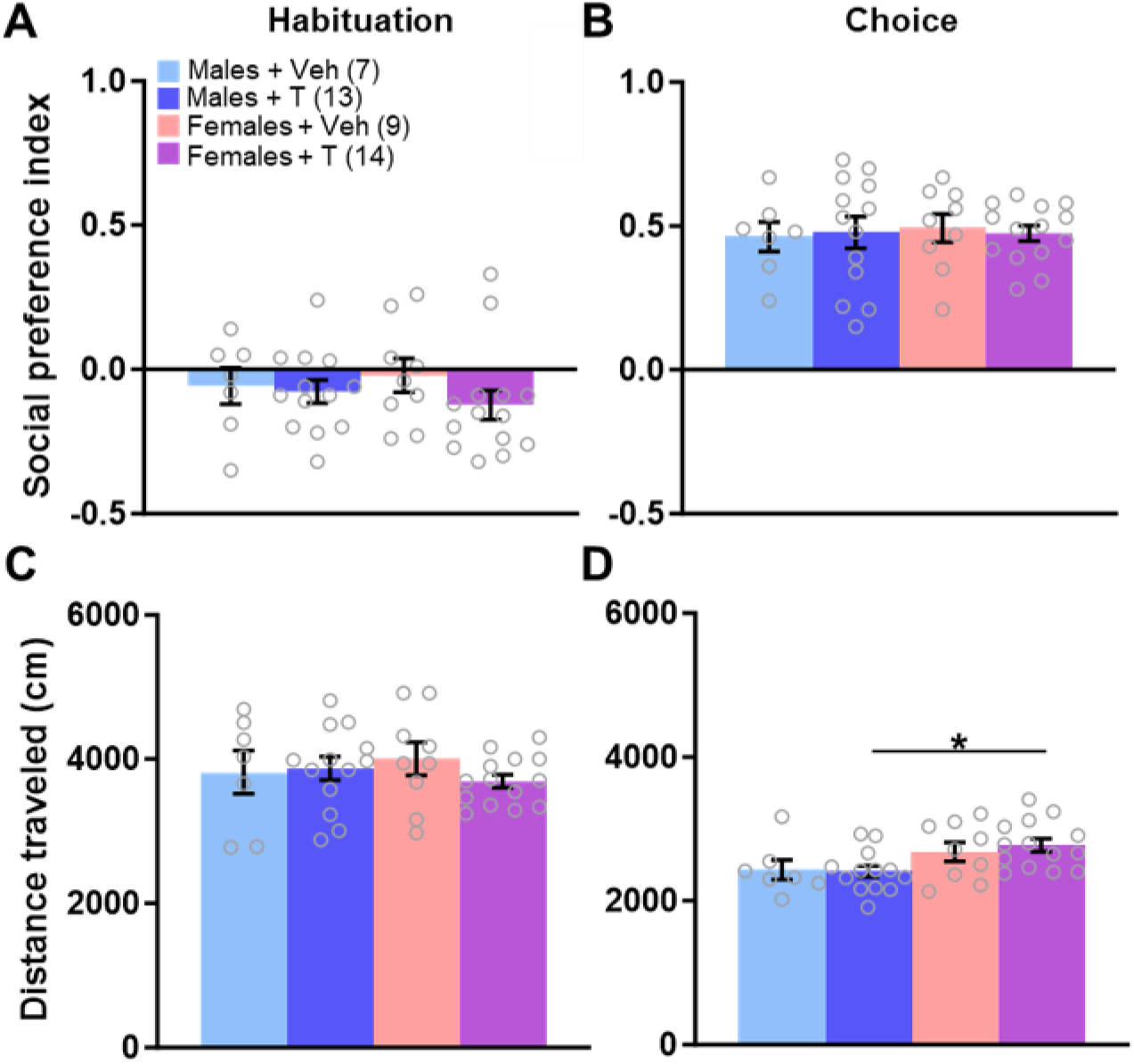
Testosterone administered on PN18 does not affect social approach behavior. (A) Male and female adolescent mice exhibited no significant differences in PI during habituation regardless of treatment with veh or T on PN18. (B) Testosterone treatment on PN18 had no effect on PI during the choice phase. (C) During habituation, all experimental groups traveled similar distances. (D) Male + Veh animals traveled less than Female + T animals during the choice phase. *p<0.05. Bars indicate mean ± SEM.

For distance traveled in the habituation phase, a two-way ANOVA revealed no main effect of sex (F_(1,39)_=0.153, p=0.698, ƞ^2^=0.004), treatment (F_(1,39)_=0.089, p=0.767, ƞ^2^=0.002), or sex x treatment interaction (F_(1,39)_=0.345, p=0.561, ƞ^2^=0.009; **Fig. 5C**). A two-way ANOVA of the distance traveled in the choice phase uncovered a significant main effect of sex (F_(1,39)_=8.207, p=0.007, ƞ^2^=0.167), no main effect of treatment (F_(1,39)_=0.105, p=0.019, ƞ^2^=0.002), and no sex x treatment interaction (F_(1,39)_=0.286, p=0.596, ƞ^2^=0.005). A Tukey post hoc test indicated that males treated on PN18 with testosterone (Males + T) had significantly lower distance traveled than Females + T (p=0.038; **Fig. 5D**), but there were no other group differences.

### A single testosterone treatment on PN18 does not induce contextual fear conditioning deficits in adults

Similarly, a two-way ANOVA of percent time spent freezing during the baseline period demonstrated a trend toward a main effect of sex (F_(1,47)_=3.887, p=0.055, ƞ^2^=0.074), no main effect of treatment (F_(1,47)_=0.779, p=0.382, ƞ^2^=0.015), and no significant phase x sex interaction (F_(1,47)_=0.081, p=0.778, ƞ^2^=0.031; **Fig. 6A**). Twenty-four hours later, freezing percentages were similar across all groups. A two-way ANOVA showed no significant main effect of sex (F_(1,47)_=2.513, p=0.120, ƞ^2^=0.049) or treatment (F_(1,47)_=0.605, p=0.441, ƞ^2^=0.012), and no interaction (F_(1,47)_=1.421, p=0.239, ƞ^2^=0.028; **Fig. 6B**).

**Figure 6.**
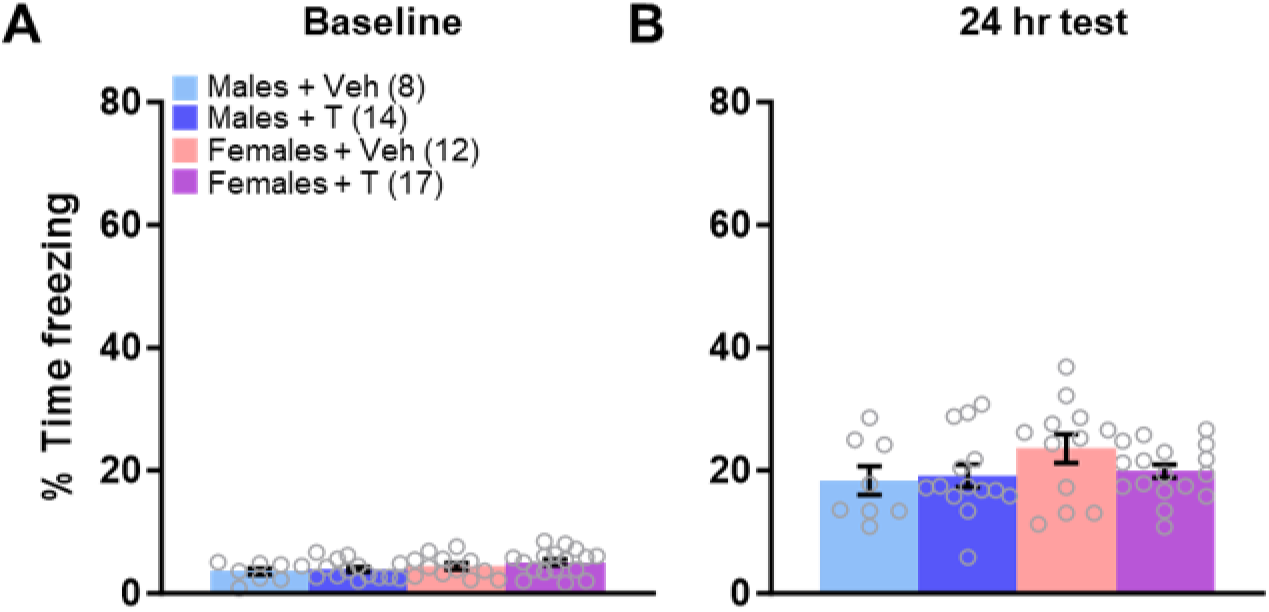
Testosterone administration on PN18 does not affect contextual fear conditioning in adults. (A) Treatment with T on PN18 did not affect percent of time spent freezing during the baseline measure. (B) There were no group differences in freezing during the 24 hr test of contextual fear memory. Bars indicate mean ± SEM.

## Discussion

Our data show that a single testosterone treatment on the day of birth results in both social deficits in juvenile, and fear memory deficits in adult, male wild-type (C57BL/6J) mice but has no effect on females. The same single s.c. injection of testosterone propionate (100 µg) (Ghahramani et al., 2014; Goel and Bale, 2013; Hisasue et al., 2010; Nakachi et al., 2015; Seney et al., 2012) administered on postnatal day 18 has no effect on social or fear memory behavior at PN30 or PN60, respectively. Considering that male rodents exhibit peak levels of testosterone several days before and after birth, while females experience negligible exposure to gonadal hormones during early development, postnatal day 18 was chosen as a timepoint outside the endogenous testosterone activity (Davis et al., 1996; McCarthy et al., 2017).

Therefore, our results indicate that excess testosterone during development is not universally detrimental but there is a sensitive period in which excess testosterone can disrupt sex-specific neural circuits and even a brief dysregulation of testosterone levels can induce long-lasting effects. Additionally, these findings indicate that both social approach behavior and fear memory exhibit sex differences in vulnerability in early development, which has significant implications for neurodevelopmental and neuropsychiatric disorders that affect males and females differently in terms of prevalence, presentation, or progression. Importantly, the observed deficits were not due to changes in body weight or motor activity that may affect general function or indicate global disruptions in physical development. The deficits were also not attributed to heightened anxiety-like behavior, which can co-occur with social deficits (Allsop et al., 2014; Felix-Ortiz et al., 2016). Notably, neonatal testosterone treatment did not affect mortality; litter size was 6.735 ± 0.392 for veh and 5.970 ± 0.388 for T, p=0.1700. Additionally, veh-treated mice exhibited behavior statistically equivalent to historical data of ours with mice that were undisturbed as neonates (no neonatal injection); both social preference indices at ∼PN30 and 24 hr fear memory results at ∼PN60 were comparable between mice administered veh and never-injected control mice we have collected over various experiments (Ferri et al., 2021, 2020, 2015; Schoch et al., 2017).

A crucial next step is to determine the mechanisms by which excess testosterone during a critical period of brain organization induces male-specific social and fear deficits. Early in development, genes on the Y chromosome orchestrate the production of testosterone by the testis in males. Testosterone can then bind directly to or be metabolized into dihydrotestosterone, which binds to, androgen receptors, or it can be aromatized to estradiol and bind to estrogen receptors. Both processes are important for distinct components of brain masculinization and defeminization (Gillies and McArthur, 2010b; McCarthy, 2011). It will be necessary to determine which or if both pathway(s) are disrupted to cause sex-specific social and fear deficits. Another critical mechanistic question concerns brain regions that may be dysregulated by excess neonatal testosterone. The medial prefrontal cortex (mPFC), amygdala, and hippocampus play important roles in both social and fear memory behavior and may be disrupted by excess testosterone early in development (Bickart et al., 2014; Bicks et al., 2015; Felix-Ortiz and Tye, 2014; Jasnow et al., 2013; Kietzman and Gourley, 2023; Kim et al., 2016; Kim and Jung, 2006; LaLumiere, 2014; Maren, 2001; Marschner et al., 2008; Zaki et al., 2022). The ventral tegmental area (VTA), nucleus accumbens (NAc), and cerebellum have also been implicated in the regulation of social behavior (Carta et al., 2019; Gunaydin et al., 2014; Musardo et al., 2022; Porcelli et al., 2019; Solié et al., 2022). Numerous sex differences in morphology and function in these brain regions have been documented, including in area volume, cell number, size, and structure, and most express androgen and estrogen receptors (Premachandran et al., 2020). Finally, fetal/neonatal testosterone during the critical organizational period of brain development has important effects on a number of downstream processes. Neurotransmitter levels, receptor expression, neuropeptide signaling, neurogenesis, synaptic programming, and cell differentiation, migration, and death, are influenced by gonadal hormones during development and may be involved in the social and fear memory deficits (Baron-Cohen et al., 2011, 2005; Ferri et al., 2018; Sara M. Schaafsma et al., 2017). Investigating these potential mechanisms will provide insight into developmental processes involved in impairments associated with neurodevelopmental and other disorders.

While we used a single, moderate dose of testosterone in this study, it will be important to determine the effects of different amounts of testosterone in future studies. This dose and timing was chosen because it has been shown to masculinize a female rodent brain (Goel and Bale, 2013; Hisasue et al., 2010; Nakachi et al., 2015; Seney et al., 2012), but the effects on the male brain and behavior have not been described. Several studies have used excess testosterone exposure *in utero;* one found that males, but not females exhibited increased density, instability, and abnormal morphology in dendritic spines of the frontal cortex, and another showed male-specific decreases in corticosterone response following restraint stress (Hatanaka et al., 2015; Wilson et al., 2020). One study administering 10x the dose used here, administered on PN2, and using RNA from the entire PN6 brain, identified 319 genes that were differentially expressed between veh and T-treated males, and decreases in estrogen receptor- and androgen receptor-responsive gene elements were found in the flanking regions of a number of those genes. In addition, levels of *Esr2* (estrogen receptor β), and *Cyp19a1* (aromatase, the enzyme that converts testosterone to estradiol) were not statistically different in males treated with veh and males treated with T, but *Esr1* (estrogen receptor α; ERα) was downregulated in T-treated male brains. *Esr1* was not different between veh- and T-treated females, however (Nakachi et al., 2015). ERα-knockout mice exhibit social deficits and a single nucleotide polymorphism (SNP) was reported to correlate with severity in social interaction deficits in a group of children with autism, although ESR1 is not considered a high confidence risk gene for ASD (Doi et al., 2018; Enriquez et al., 2021; Ervin et al., 2015). Therefore, the possibility of ESR1-mediated mechanism of approach and avoidance deficits will be important to examine in our paradigm. Relatedly, investigation of additional timepoints between PN0 and PN18 will be valuable to help pinpoint periods of vulnerability, and while here we treated the entire litter with veh or testosterone and those same-treatment mice remained housed together, future studies will address whether mixed-treatment mice housed together exhibit social and fear memory deficits similar to those described here.

Importantly, complications of pregnancy including PCOS, pre-eclampsia, and gestational diabetes that result in increased risk of neurodevelopmental disorders in the offspring involve a number of complex factors in addition to increased levels of testosterone. Likewise, an interaction of genes or environmental insults and sex hormone levels likely contribute to the development of neuropsychiatric conditions, and there is some evidence of this (Baron-Cohen et al., 2015; Bordt et al., 2024; Sara M Schaafsma et al., 2017; Schaafsma and Pfaff, 2014; Young and Pfaff, 2014). However, we have shown that excess testosterone in early development alone is sufficient to induce neurodevelopmental deficits in mice, which paves the way for future studies. Obvious ethical constraints prohibit well-controlled manipulations in humans, and it is not possible to safely study testosterone levels in a human fetus mid-gestation during the critical period of brain masculinization. Relatedly, the relationship of blood or brain levels of testosterone and more accessible samples like maternal blood levels or those measured during amniocentesis, which is usually only indicated in high-risk pregnancies, is not known. These studies are valuable, but are also complicated by heterogeneity in human subjects, lack of information, small sample size, and findings that fail to replicate. Animal models can help identify possible biomarkers that are modifiable as possible treatment targets, as there are several pharmacological agents in use that modulate steroid hormone levels. In conclusion, while we do not present the manipulation in this study as a model of any specific disorder, we propose that it is a valuable paradigm in which dysregulation of sex-specific and time-sensitive developmental pathways can be used to investigate differential vulnerabilities to behavioral deficits, which may be relevant to a number of neuropsychiatric conditions that express sex differences.

## Acknowledgements

This work was supported by the National Institutes of Health (K01 MH119540) and University of Iowa Hawkeye Intellectual and Developmental Disabilities Research Center (Hawk-IDDRC; NICHD; P50 HD103556; PI Abel and Strathearn). We would like to thank Dr. Shane Heiney and the Neural Circuits and Behavior Core at the University of Iowa and Emily Hagan, Danielle Preuschl, and Charlotte Tesar for their input and support.

## Author contributions

SLF designed the experiments, performed the experiments, analyzed the data, and wrote the manuscript. PQL performed the experiments, analyzed the data, and wrote the manuscript. KLB and MRB performed the experiments and analyzed the data.

## Additional Information

The authors declare no competing interests.

